# Whole-genome DNA methylomes of Tree shrew brains reveal conserved and divergent roles of DNA methylation on sex chromosome regulation

**DOI:** 10.1101/2024.06.05.597676

**Authors:** Dongmin Son, Yifan Kong, Yulian Tan, Ting Hu, Lei Shi, Soojin V. Yi

**Author notes:** Correspondence: Lei Shi, Soojin Yi.

## Abstract

The tree shrew (*Tupaia belangeri*) is a promising emerging model organism in biomedical studies, notably due to their evolutionary proximity to primates. To enhance our understanding of how DNA methylation is implicated in regulation of gene expression and the X chromosome inactivation (XCI) in tree shrew brains, here we present their first genome-wide, single-base-resolution methylomes integrated with transcriptomes from prefrontal cortices. We discovered both divergent and conserved features of tree shrew DNA methylation compared to that of other mammals. DNA methylation levels of promoter and gene body regions are negatively correlated with gene expression, consistent with patterns in other mammalian brains studied. Comparing DNA methylation patterns of the female and male X chromosomes, we observed a clear and significant global reduction (hypomethylation) of DNA methylation across the entire X chromosome in females. Our data suggests that the female X hypomethylation does not directly contribute to the gene silencing of the inactivated X chromosome nor does it significantly drive sex-specific gene expression of tree shrews. However, we identified a putative regulatory region in the 5’ end of the X inactive specific transcript (*Xist)* gene, a key gene for XCI, whose pattern of differential DNA methylation strongly relate to its differential expression between male and female tree shrews. We show that differential methylation of this region is conserved across different species. Moreover, we provide evidence suggesting that the observed difference between human and tree shrew X-linked promoter methylation is associated with the difference in genomic CpG contents. Our study offers novel information on genomic DNA methylation of tree shrews, as well as insights into the evolution of X chromosome regulation in mammals.

## Introduction

The tree shrew (*Tupaia belangeri*) is a small mammal widely found in Southeast Asia and China. The tree shrew offers several advantages to be used as a model species for biomedical studies. For example, the tree shrew has a small body size and short life span, making it easy to rear in laboratories for experimental studies (Yao, 2017). On the other hand, the tree shrew has a high brain to body ratio, and exhibit several conditions that can model human disorders (Li et al., 2024). Several studies have demonstrated greater genetic similarities between tree shrew and primates than between rodents and primates, especially in genes associated with neuropsychiatric disorders and infectious diseases (Fan et al., 2013; Yamashita et al., 2012). Indeed, the tree shrew is the closest group of mammals to primates, providing a useful model system especially in neuroscience (Fan et al., 2013).

While genetic and transcriptomic studies of tree shrews are becoming available in the literature, epigenetic mechanisms such as DNA methylation of tree shrews have been little explored so far. Given the important role of DNA methylation in regulatory processes such as regulation of gene expression, neuropsychiatric diseases and the X chromosome inactivation (XCI), such data will advance our understanding of regulatory evolution and enhance the utility of tree shrew as a model species. Here, we generated and analyzed whole genome DNA methylation maps (methylomes) of tree shrew prefrontal cortex from three males and females. Integrating them with transcriptomic data, we demonstrate genome-wide influence of DNA methylation on gene expression, including the presence of CG and CH methylation which are both associated with gene expression.

Moreover, we examined the role of DNA methylation in the regulation of X chromosome inactivation (XCI). Notably, we show that global differential DNA methylation between the male and female X chromosome is not a driver of XCI in tree shrews. On the other hand, we newly annotated the X inactive specific transcript (*Xist)* gene and identified an evolutionary conserved locus of regulation in the 5’ end of the gene. Comparing patterns of DNA methylation and X chromosome regulation across different species, we show that the evolutionary patterns of X chromosome DNA methylation is closely associated with the difference in genomic CpG contents. Additionally, we identified putatively Y-linked genomic segments and their hypomethylation. These novel findings illuminate conserved and divergent patterns of genomic DNA methylation and regulation of X chromosome in mammals.

## Results

### Genomic DNA methylation in tree shrew

We generated whole-genome bisulfite sequencing (WGBS) data from the prefrontal cortex (herein referred to as ‘PFC’) of 6 Chinese tree shrews (3 males and 3 females) to produce DNA methylomes at nucleotide resolution (Table S1). This marked the first whole genome methylome study of the tree shrew, integrated with transcriptome data. Previous studies of human and mouse brain methylomes have identified CG and non-CG (CH where H is A, T, C) DNA methylation (Jeong et al., 2021; Lister et al., 2009; Zeng et al., 2012). Tree shrew PFCs were also highly methylated at CG sites while CH methylations were observed at lower levels compared to CG methylation (Fig.1A). We observed significantly lower levels of CG methylation in promoter regions (defined as 2kb upstream of the TSS) than in gene bodies and a sharp drop in CG methylation near transcriptional start sites (TSS) (Fig. 1B, Fig. S1). Gene body CG methylation levels were higher than nearby intergenic regions. DNA methylation at non-CG sites remained relatively consistent in all genomic context with a slight drop near TSS (Fig. S2). These patterns were similar across all chromosomes except for the chromosomes 13 and 26, where chromosome 26 showed a significant drop near the end of genes and chromosome 13 exhibited strikingly lower methylation levels across all genomic contexts (Fig. S2).

**Figure 1.**
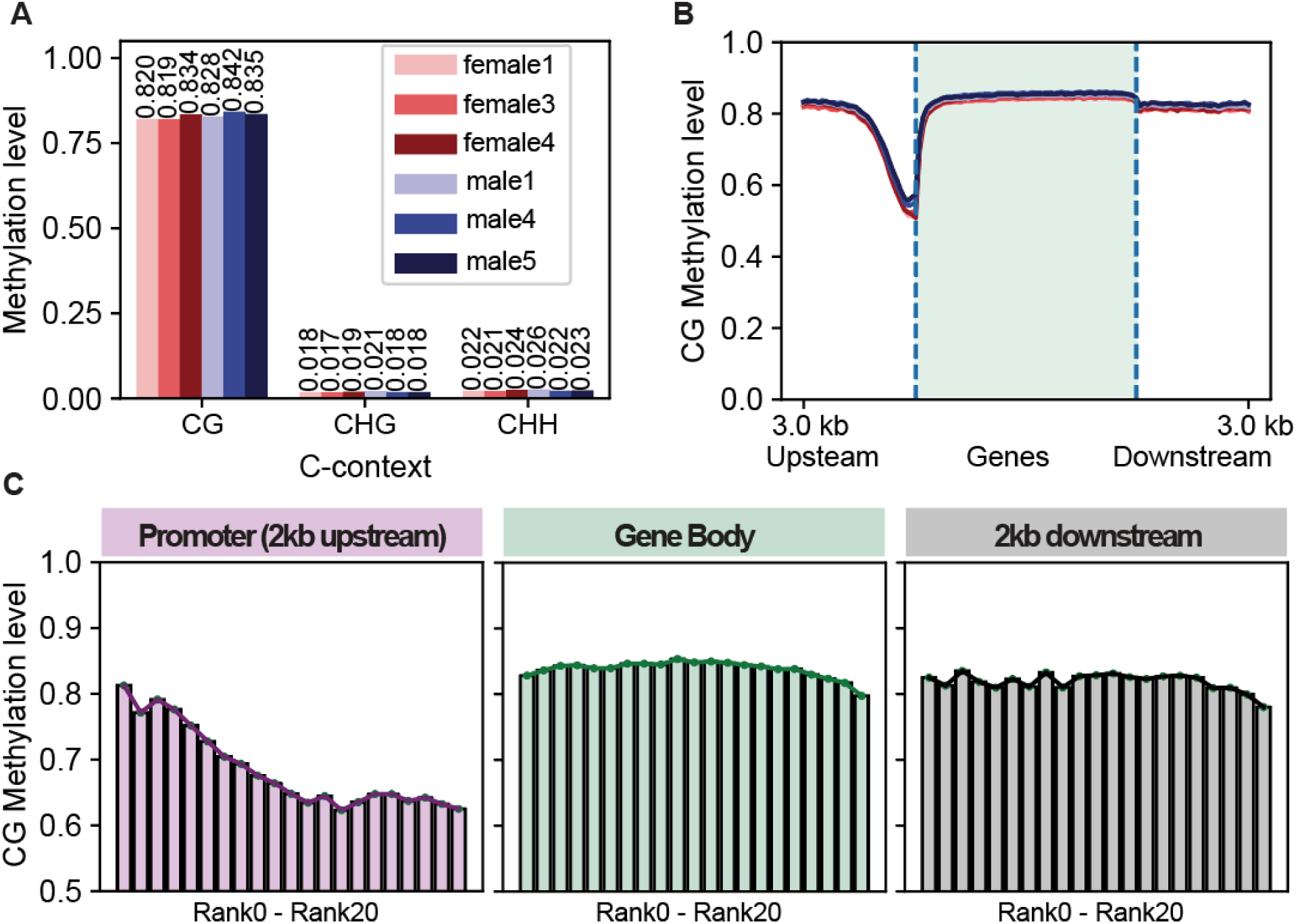
DNA methylomes of tree shrew and the correlation between DNA methylation and gene expression. Chromosomes 13 and 26 (1,001 genes) were excluded due to their unique patterns compared to other chromosomes. (A) Global (weighted) mean DNA methylation levels of CG, CHG, and CHH in each sample exhibit high levels of CpG methylation and low levels of CH methylation. (B) Mean (weighted) DNA methylation across gene bodies of 22,741 protein-coding genes in the autosomes and the X chromosome demonstrate decreases of DNA methylation near the transcription start sites. (C) Mean (weighted) DNA methylation of promoter, gene body, and intergenic regions in 20 groups of genes with different expression levels, ranging from rank 0 to rank20 (higher expressed genes from left to right). A negative correlation is observed in promoters, while a bell-shaped correlation is observed in gene bodies.

Integrating methylomes with RNA-seq data from the same samples, all 22,741 protein-coding genes were ranked into 0-20 (lowest to highest) according to their expression levels (Fig.1C). While genes in all ranks showed hypomethylation near the TSS, highly expressed genes had the lowest methylation levels near TSSs while 0 or lowly expressed genes had relatively higher methylation levels (Fig. 1C, Fig. S1B), resulting in a significant negative correlation between gene expression and promoter methylation (Fig. S4, Table 1), even though for gene bodies, it was only significant for protein coding genes, not in lncRNA genes (Table 1). In addition, protein-coding genes showed a more pronounced methylation drop near TSS compared to lncRNA genes (Fig. S2). This discrepancy may be partly attributed to incomplete annotations of some lncRNA genes. In the gene bodies, the correlation was bell-shaped similar to previous findings in humans (Jjingo et al., 2012), with lowly expressed genes and highly expressed genes both having relatively lower methylation compared to the median expressed genes (Fig.1C, Fig. S4).

**Table 1.** Correlation analysis of mean promoter and gene body DNA methylation levels and ranked gene expression of genes of an autosome and the X chromosome of the tree shrew. (**A**) for protein-coding genes and (**B**) for lncRNA genes. Spearman’s rank correlation coefficeint (ρ) and p-value for each chromosome in males and females are shown.

| **(A)**<br>**Protein-Coding** | Chr | Number of genes analyzed | Genomic Region | Spearman's $\rho$ (p-value) (Male) | Spearman's $\rho$ (p-value) (Female) |
| --- | --- | --- | --- | --- | --- |
| | Chr 8 | 597 | Promoter | -0.31 ($7.24 \times 10^{-15}$) | -0.34 ($4.95 \times 10^{-17}$) |
| | | | Gene body | -0.29 ($5.49 \times 10^{-13}$) | -0.34 ($2.31 \times 10^{-17}$) |
| | Chr X | 823 | Promoter | -0.34 ($5.58 \times 10^{-23}$) | -0.31 ($6.34 \times 10^{-20}$) |
| | | | Gene body | -0.31 ($1.42 \times 10^{-18}$) | -0.26 ($2.83 \times 10^{-13}$) |
| **(B)**<br>**lncRNA** | Chromosome | Number of genes analyzed | Genomic Region | Spearman's $\rho$ (p-value) (Male) | Spearman's $\rho$ (p-value) (Female) |
| | Chr 8 | 1759 | Promoter | -0.06 ($1.76 \times 10^{-2}$) | -0.04 ($7.96 \times 10^{-2}$) |
| | | | Gene body | -0.07 ($1.73 \times 10^{-3}$) | -0.10 ($1.53 \times 10^{-5}$) |
| | Chr X | 1058 | Promoter | -0.09 ($1.14 \times 10^{-3}$) | -0.11 ($2.60 \times 10^{-4}$) |
| | | | Gene body | -0.04 ($1.79 \times 10^{-1}$) | -0.04 ($2.19 \times 10^{-1}$) |

### Differential DNA methylation of female and male tree shrew X chromosomes

We compared DNA methylation of autosomes and the X chromosome in the tree shrews PFC. We selected chromosome 8 as a representative autosome for comparison with the X chromosome, since it has a similar size and CG-content to the X chromosome. The mean coverage depth at CpG sites across the X chromosome was approximately twice as high in females as in males, as expected from the 2:1 ratio of the X chromosome in females compared to in males (Fig. S3). We identified a region on the X chromosome where reads were mapped only in males, both in the methylomes and in the transcriptomes (Fig. S3). We inferred that this included a part of the Y-linked regions that were incorrectly assembled into the X chromosome in the reference assembly (Chr X: 4542400-6144400) and subsequently removed it from further analysis (Methods).

We observed that the X chromosomes of females displayed significantly lower levels of DNA methylation in gene bodies and promoters compared to those of males (*P* = 6.13 × 10^-110^ and 1. 02 × 10^-195^ for promoters and gene bodies, respectively, Mann-Whitney U test, Fig. 2D, Table 2). Comparisons to the autosomes demonstrated that this pattern was due to the reduced DNA methylation in females, or ‘female hypomethylation’ (Fig. 2A and B, Fig. S4). This pattern was consistently observed across the entire X chromosome and was pervasive across different functional regions, and also in non-CpG methylation (Fig. 2C, Fig. S6). The observation of female X hypomethylation was not due to a bias from different read depths between females and males, as we observed the same patterns when we re-analyzed equal depth data by randomly sampling half of the sequencing reads on female X chromosomes (Fig. S7). The observed pattern resembled what was observed in marsupial koalas and differed from that in humans, where promoters were typically hypermethylated in females (Singh et al., 2021).

**Figure 2.**
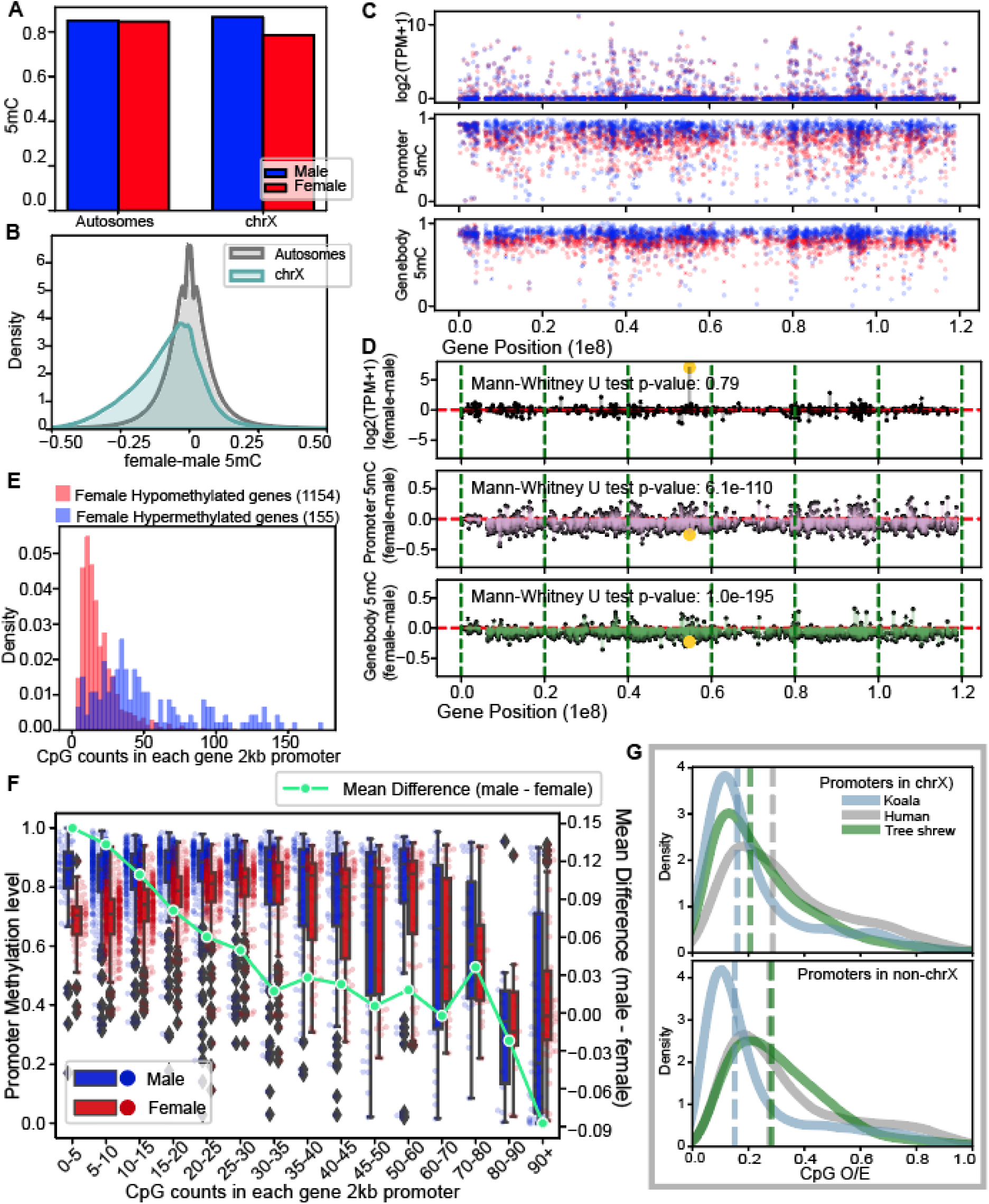
Global patterns of female X hypomethylation in the tree shrew PFC and its association with CpG counts. (A) Mean fractional DNA methylation levels of all CpGs in males and females demonstrates that the female X chromosome is globally hypomethylated compared to chromosome 8 and the male X chromosome. (B) Distributions of DNA methylation level differences of CpG sites between females and males in autosomes and the X chromosomes show that the X chromosome is generally hypomethylated. (C) The distribution of expression levels, promoter DNA methylation levels, and gene body DNA methylation levels of genes (1857 genes including both protein-coding genes and lncRNA genes) across the X chromosome in females (red) and males (blue). (D) The differences between females and males for expression, promoter methylation and gene body DNA methylation of genes across the X chromosome. The yellow dot in the gene expression plot represents the *Xist* gene, which is up-regulated in females. (E) Genes with female hypomethylated (mean 5mC male-female > 0.05, 1154 genes) promoters tend to have fewer CpGs compared to those with female hypermethylated (mean 5mC male-female < –0.05, 155 genes) promoters. (F) Comparisons of DNA methylation (Y-axis on the left) levels and their differences between males and females (Y-axis on the right) according to the numbers of CpGs in promoters (X-axis). Female promoters are clearly hypomethylated compared to male promoters when CpG counts are low. As CpG counts increases, both promoters are generally lowly methylated. Promoters with large CpG counts (>80) are on average female hypermethylated. (G) A comparison of promoter CpG O/E across three species (human, tree shrew and koala) demonstrates the similarity between the koala and the tree shrew compared to the human, specifically in the X-linked promoters.

**Table 2.** Differences in DNA methylation levels and expression levels between female and male tree shrews, along with Mann-Whitney U test results and its p-values.

|  |  | Number of Genes compared | Mann-Whitney U test p-value |
| --- | --- | --- | --- |
| Chr8 | Promoter 5mC | 2324 | 0.12 |
|  | Gene body 5mC | 2245 | 0.02 |
|  | Expression | 2341 | 0.86 |
| Chr X | Promoter 5mC | 1790 | $6.13 \times 10^{-110}$ |
| | Gene body 5mC | 1667 | $1.02 \times 10^{-195}$ |
|  | Expression | 1856 | 0.79 |

### Promoter methylation difference between females and males associate with CpG counts

We found that the degree of tree shrew female hypomethylation compared to males in promoters was highly depended on the density of CpGs within the promoter regions. First, female hypomethylated promoters had fewer numbers of CpGs compared to female hypermethylated genes (Mann-Whitney U test, P = 6.70 × 10^-39^, Fig. 2E, note that these counts reflect CpG density as the total number of nucleotides are the same for all 2kb-sized promoters). Second, the difference between female and male promoter methylation decreased as the number of CpGs with promoters increased (Fig. 2F).

We propose that the dependency of promoter hypomethylation on CpG counts can explain the observed difference between species. We compared the distribution of CpG contents in human, using the metric CpG Observed/Expected ratio (CpG O/E) (Elango & Yi, 2008). We found that while human X chromosome and autosomes included a large number of high CpG O/E promoters, koala X promoters mostly consist of low CpG O/E promoters (Fig. 2G, Fig. S8). Tree shrew promoters resembled the pattern observed in koalas, where most X-linked promoters had low CpG O/E (Fig. 2G, Fig. S8). In contrast, CpG O/E from autosomes were similar between humans and tree shrews compared to that in koalas (Fig. S8).

### Hypomethylation of the tree shrew female X does not drive sex-specific expression

If the primary functional outcome of X chromosome DNA hypomethylation is an up-regulation in gene expression, as supported by the negative correlation observed between DNA methylation levels and gene expression, then we would expect to see an overall increase in gene expression across female X-linked genes. However, this was not the case and we observed no global difference of gene expression between the male and female X chromosomes, as expected under the functional XCI (Fig. 2D, Table 2). Similarly, chromosome 8, where there was no discernible DNA methylation difference between males and females, showed no global difference of gene expression between males and females (Fig. S5, Table 2). Therefore, we concluded that female hypomethylation of the tree shrew X chromosome did not lead to up-regulation of genes.

Out of the total 32,302 autosomal genes including both protein-coding and non-coding RNA genes, 479 genes were significantly differentially expressed between males and females (Adjusted p-value < 0.1 based on DESeq2, Fig. 3A). In comparison, 14 out of 783 X-linked genes were significantly differentially expressed (Fig. 3C). The proportion of sex-specific genes was not statistically different similar between the autosomes and X chromosomes (z-score test, P = 0.48).

**Figure 3.**
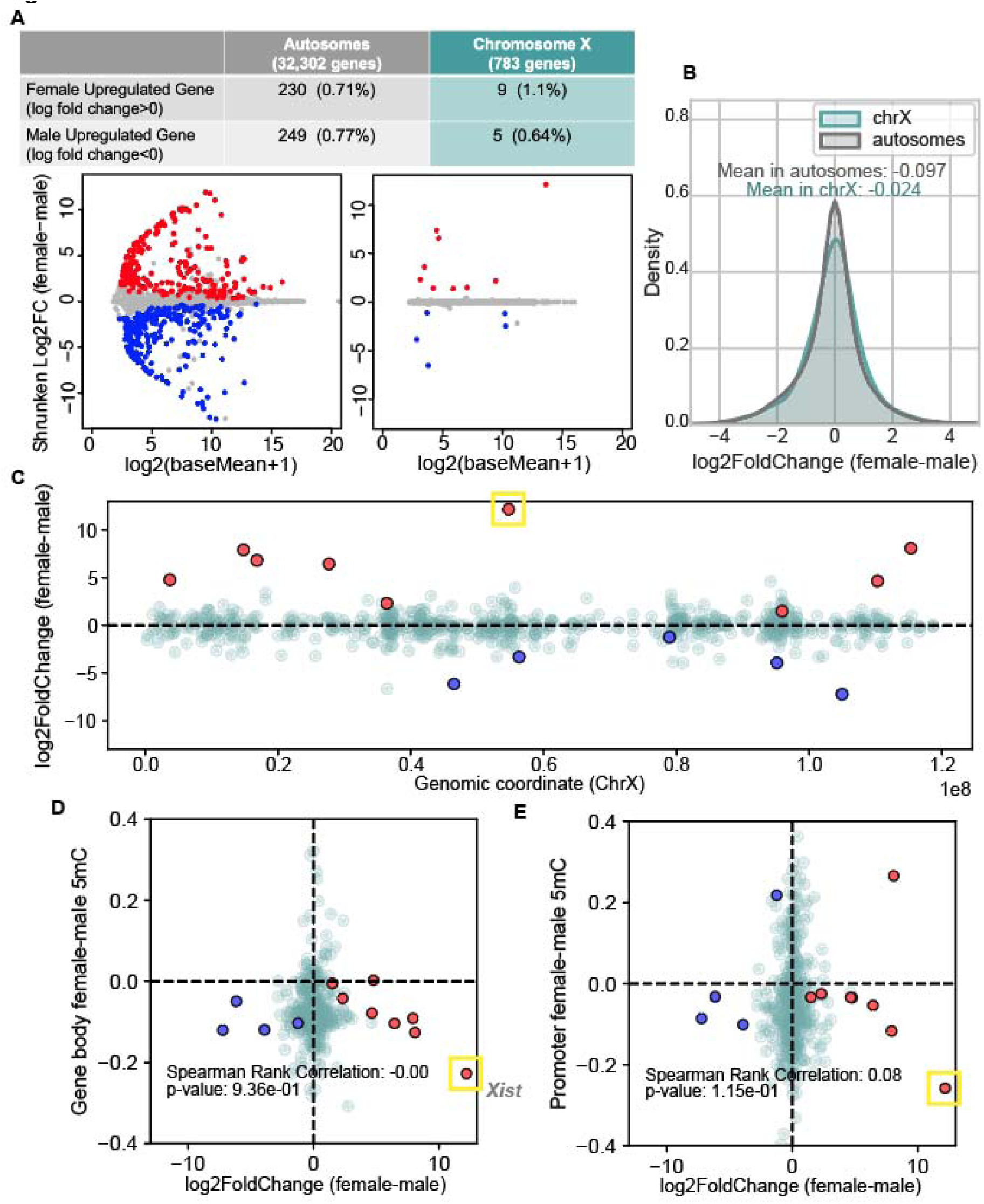
Patterns of differential expression between males and females in relation to differential DNA methylation. Sex-specific expressed genes and their correlation with DNA methylation levels. (A) The MA plot illustrating differentially expressed genes across autosomes (left) and the X chromosome (right). Ashr-shrunken log fold-change values are used for the visualization. Blue dots represents male up-regulated genes and red dots represents female up-regulated genes. (B) The density distribution graph displays the log-transformed female-to-male expression ratio for genes. There was no significant difference between the X chromosome and autosomes (Mann-Whitney U test p-value = 0.13). (C) The distribution of female (red) and male (blue) up-regulated genes identified by DESeq2 across the X chromosome. The *Xist* gene is marked with a yellow box. (D, E) The Y-axis represents the difference in mean DNA methylation levels between females and males is across gene bodies (D) and promoters (E). The X-axis represents the log2 fold-change of female-to-male expression difference. Spearman’s rank correlation coefficient and p-value are reported.

Across the entire set of autosomes, there was a slight excess of male-biased genes, characterized by a greater number of male up-regulated genes compared to female up-regulated genes. This trend remained consistent when different p-value thresholds were employed (Fig. S9). On the other hand, there was an excess of female biased genes on the X chromosome (Fig. S9). The average log2 fold-change of female to male expression was –0.097 for autosomes and –0.024 for chromosome X. However, it is worth noting that this difference was not significant (Mann-Whitney U test, *P* = 0.13, Fig. 3B) and the average log2 fold-change values showed variability depending on the stringency of filtering steps in the analysis (Fig. S9).

We examined whether female chromosome X hypomethylation contribute to the sex-specific expression. Among the genes on the X chromosome, 443 gene had available information regarding log2 fold-change expression and the gene body DNA methylation level, while 439 genes had available information regarding log2 fold-change expression and the promoter DNA methylation level. Including these genes, we conducted an analysis of the relationship between gene expression and DNA methylation levels (Fig. 3D and E). We found that the methylation level difference between females and males on the X chromosome did not exhibit any significant correlation with the gene expression difference between females and males (*P* = 0.94 and *P* = 0.12 for Spearman’s rank correlation test, for gene bodies and promoters, respectively). Likewise, there was no significant trend between DNA methylation difference and gene expression difference when only significant sex-specific genes were analyzed (*P* = 0.46 and 0.57 for Spearman’s rank correlation test, for gene bodies and promoters, respectively) (Fig. 3D and E). These observations suggest that female X chromosome hypomethylation does not contribute significantly to sex-specific gene expression.

### Differential methylation of the *Xist* associated with its sex-specific expression

The *Xist* gene is known for its key role in the initiation and maintenance of XCI in mammals (Penny et al., 1996; Pontier & Gribnau, 2011). However, how *Xist* gene is regulated in the tree shrew remained unknown prior to our study. In fact, the current tree shrew genome database did not include a specifically annotated *Xist* gene. Here, we first identified the putative *Xist* gene region in the tree shrew genome. Briefly, we conducted a BLASTN search using the human *Xist* gene sequence as a query against the tree shrew reference genome to obtain a potential genomic coordinate of *Xist* gene. We then identified novel transcripts from tree shrew RNA-seq samples using StringTie’s functionality for *de novo* transcript assembly (See Method). We identified a single gene from these novel transcripts. We found that longer novel transcripts were produced from female samples compared to male samples (Fig. 4A). Furthermore, this gene exhibited extremely high expression levels in our female data (Fig. 3C). The new annotation puts the tree shrew *Xist* on chr X: 54,721,977-54,759,147.

**Figure 4.**
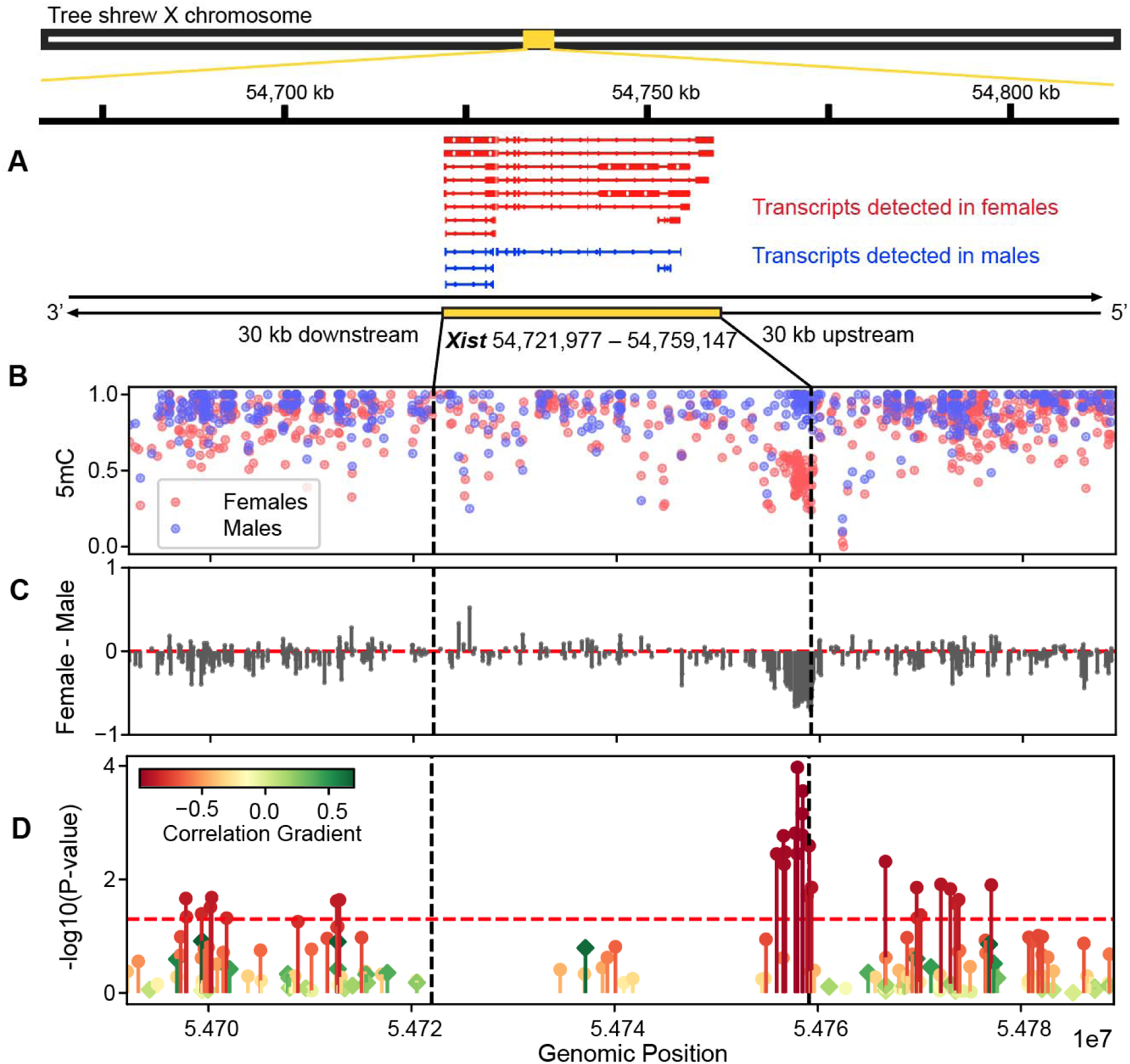
DNA methylation difference between females and males for the newly annotated tree shrew *Xist* gene. (A) Longer transcripts are detected in female samples compared to male samples in the *Xist* gene region. (B) Fractional methylation levels of CpG sites around *Xist* are indicated and (C) the differences of DNA methylation levels of each CpG site between females and males are calculated. A CpG island near the 5’ end of the gene displays marked female hypomethylation. (D) The methylation levels at each CpG site are correlated with gene expression levels using six samples. The Y-axis values represent the significance of the Pearson correlation, with a red horizontal line indicating the threshold for a p-value of 0.05. The color gradients indicate their corresponding Pearson correlation coefficient, with red for a negative correlation and green for a positive correlation). Circular points represent female-hypomethylated CpGs, while diamond-shaped points indicate male-hypomethylated CpGs.

The *Xist* displays the most pronounced differential expression between the male and female tree shrew PFC (Fig. 2D, Fig. 3C and D, indicated by a yellow box), consistent with its key role in X-chromosome inactivation (XCI) in eutherian mammals. While *Xist* is a rapidly evolving non-coding RNA, specific regions are known to exhibit a high degree of conservation among eutherian mammals. Notably, the 5’ end of *Xist* is known to be relatively well conserved compared to other regions (Brown et al., 1992; Hendrich et al., 1993; Rauch et al., 2009). Specifically, the 5’ region of *Xist*, including a CpG island, is hypomethylated on the inactive X chromosome and highly methylated on the active X chromosome to maintain *Xist* repression in human and mouse (Hendrich et al., 1993). DNA methylation in this region is known to play an active role in controlling *Xist* transcription in mouse (Beard et al., 1995; Norris et al., 1994; Panning & Jaenisch, 1996) and in human somatic cells (Tinker & Brown, 1998). We therefore investigated differential DNA methylation between females and male tree shrews within *Xist* region (across the 37.2kb gene body and 30kb upstream and downstream regions) and identified a CpG island near the 5’ end that exhibited a striking pattern of female hypomethylation (Fig. 4B, C). We examined the relationships between DNA methylation of CpGs in this region and the *Xist* gene expression levels across the six samples (Fig. 4D) and found several differentially methylated CpG positions within this female hypomethylated CpG island exhibiting significant correlations (13 CpGs located near the 5’ end of *Xist* gene with Pearson correlation, *P* < 0.05). These results indicate that the *Xist* gene is regulated by methylation of CpGs that are female-hypomethylated and inversely correlating with expression, resulting in its active expression in female. Our results suggest that the 5’ end CpG island and its hypomethylation in the inactive X chromosome are conserved across mouse, human, and tree shrew. We processed the public WGBS data of these species and found it can be consistently detected using WGBS data (Fig. S10).

### Identification of Y-linked contigs and the lower methylation level of the Y chromosome

Leveraging the availability of both female (XX) and male (XY) samples, we explored the level of DNA methylation in the Y-chromosome, a topic sparsely explored so far. The reference tree shrew genome lacks the Y chromosome assembly. To detect the Y-linked segments, we developed a method to compare CpG read-depths between female and male samples. Scanning the contigs currently unassembled using this method, we detected potentially Y-linked segments as those where female samples showed a lack of read counts compared to male samples. We demonstrate our methodology by computing 1 – (mean read-depth in females)/(mean read-depth in males) in Figure 5. For autosomal-linked contigs, since mean read-depths should be similar between males and females, this metric should be near zero. For X-linked contigs, the mean read-depth in females should be greater than that in males, and this metric should be less than zero. For Y-linked contigs, this metric should be closer to 1. Indeed, we observed three distinct peaks as expected: the left peak likely represents X-linked contigs, the middle peak corresponds to autosome-linked contigs, and the right peak indicates Y-linked contigs (Fig. 5A).

**Figure 5.**
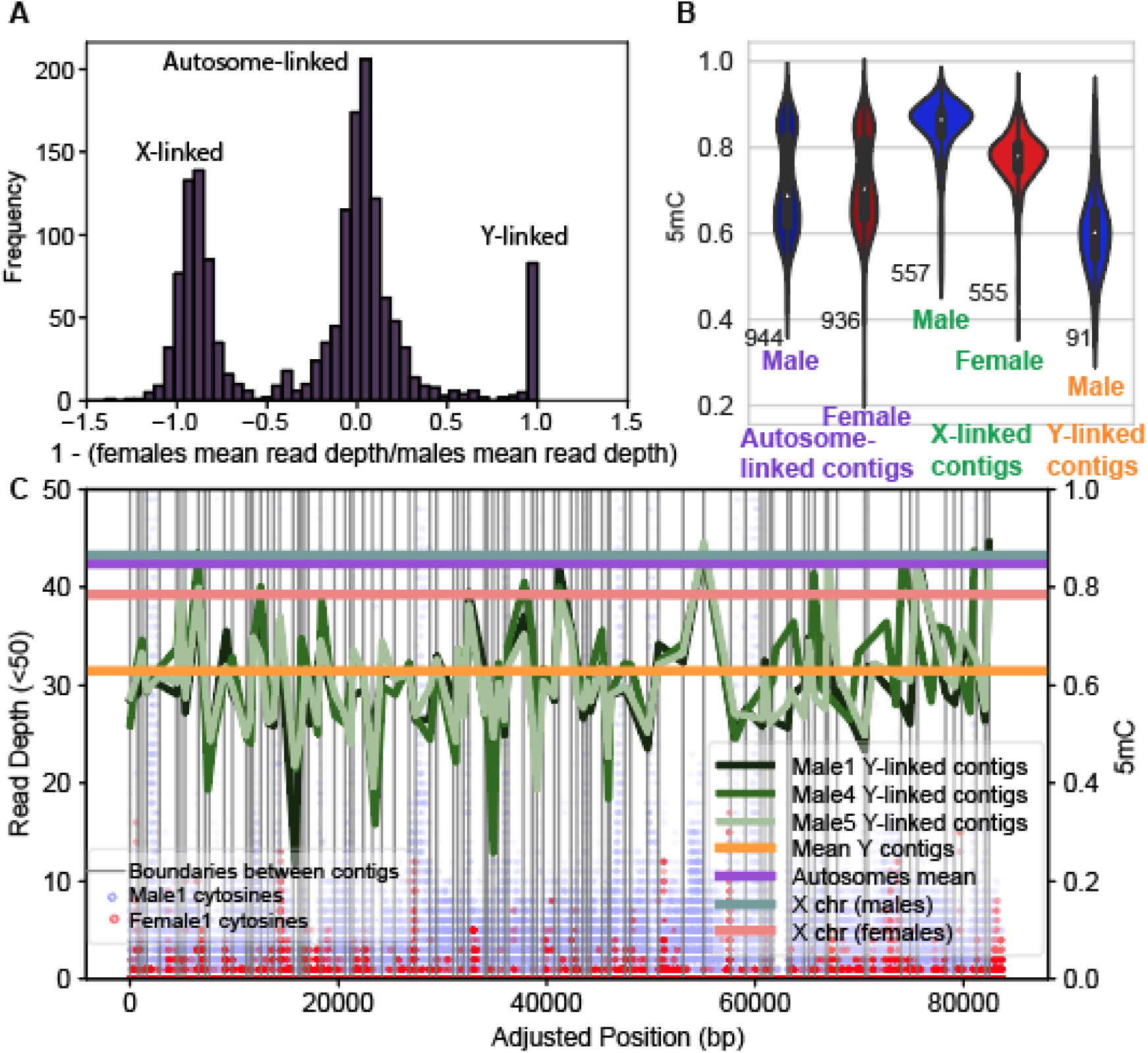
DNA methylation in the Y-linked contigs of the tree shrew PFC. (A) The distribution of the metric [1-(mean read-depth in females)/(mean read-depth in males)] for contigs. Three peaks are observed, which potentially correspond to from X-linked, autosome-linked, and Y-linked contigs. (B) Mean methylation levels for contigs in each peak. Notably, putatively X-linked contigs displayed female X hypomethylation. The numbers of contigs included in the sets are indicated. (C) (Left axis) The read-depth of cytosines in the CG context are visualized for male 1 and female 1 sample across each Y-linked contig. The contigs are sorted and connected create an adjusted position. (Right axis) The methylation levels of these contigs, compared with autosomes and the X chromosome. The Y-linked contigs exhibit markedly lower methylation levels than other chromosomes.

We applied a filter for values greater than 0.75 to identify a list of 93 potential Y-linked contigs. The read-depth of each cytosine across these contigs was graphically represented to show a lack of female counts mapped (Fig. 5C). Subsequently, we calculated and compared the methylation levels of these contigs against those of other chromosomes. Notably, the Y chromosome in the tree shrew PFC exhibited markedly lower methylation levels compared to both autosomes and the X chromosome (Fig. 5B, C). Moreover, contig18904 (23,877 bp in size) is identified as Y-linked using this methodology, wherein the sex-determining region Y (*Sry*) genes of humans and rhesus monkeys align in the tree shrew genome (Fig. S11), exhibiting 79% identity for human-tree shrew alignment and 82% identity for rhesus monkey-tree shrew alignment based on BLASTN (See Method). Interestingly, no expressed *Sry* transcripts are detected in our tree shrew prefrontal cortex transcriptome data of male samples. *Sry* gene expression is known to be tissue-specific and under the control of DNA methylation in mice and humans. (Gimelli et al., 2006; Larney et al., 2014; Nishino et al., 2004). Our study reveals the putative *Sry* locus in the tree shrew genome (Fig. S11) and indicates its extremely low DNA methylation and low expression in PFCs.

## Discussion

This study presents the first genome-wide analysis of DNA methylation of the emerging model species, the tree shrew. In human and mouse, DNA methylation of cis-regulatory regions is known to dampen the expression of associated genes (Schübeler, 2015). DNA methylation of gene bodies is also known to contribute to regulation of gene expression, although the directionality is not as straightforward (Jjingo et al., 2012; Schübeler, 2015). We observed significant and strong negative correlations between promoter DNA methylation and gene expression across the genome, which is consistent with the aforementioned model as well as with previous studies (Al Adhami et al., 2022; Jjingo et al., 2012; Mendizabal et al., 2019; Rauch et al., 2009). Gene body DNA methylation was also associated with expression, although its effect was less pronounced compared to promoter methylation.

We observed similar levels of overall gene expression between the male and female tree shrew X chromosomes, as expected under functional XCI. Interestingly, female tree shrew X chromosome exhibited substantially lower level of DNA methylation compared to the male X chromosome (Fig. 2, Fig. S4). From the comparisons to autosomes and the male X chromosome, we demonstrate that this was due to the reduction of DNA methylation in the female X chromosome in tree shrews.

Examining literature that have specifically addressed the global DNA methylation difference between the male and female X chromosome using chromosome-wide methods, Hellmann and Chess (Hellman & Chess, 2007) demonstrated that gene bodies were hypomethylated in the human female X chromosome, and Keown et al (Keown et al., 2017) showed that intergenic regions were hypomethylated in mouse female X chromosome. Sun et al. (Sun et al., 2019) and Singh et al. (2021) analyzed WGBS data and showed that human female X chromosome was hypomethylated compared to the male X chromosome except for the promoter regions. Here we show that in tree shrews, female X chromosome is globally hypomethylated compared to male X chromosome and autosomes, which is similar to the pattern observed in a marsupial, koalas (Singh et al., 2021).

While we observed that while the majority of the promoters (86.6%, Fig. 2E) was hypomethylated in female X, some promoters were hypermethylated in the female X chromosome of tree shrew. Intriguingly, these promoters were those harboring high density of CpGs (Fig. 2E). CpG-rich promoters are known to be associated with highly and broadly expressed genes in diverse vertebrate species (Elango & Yi, 2008). Our findings of the close association between female promoter hypomethylation and CpG density provide insights into the observed difference between species where similar hypomethylation was observed in the promoters of female tree shrews and koalas, in contrast to the hypermethylation detected in human female X promoters. We show that CpG O/E ratio within X chromosome promoters is relatively similar between koalas and tree shrews, while the CpG O/E ratio between humans and tree shrews is similar across other chromosomes (Fig. S8). In addition, a recent in-depth analysis of DNA methylation difference between the female and male human X chromosome demonstrated that CpG-rich promoters tend to be hypermethylated in female X, suggesting that a similar principle could be applied to variation within the human genome (Morgan et al., 2024).

It is noteworthy that we found little difference in gene expression but strong differential methylation between the male and female X chromosomes, especially considering the pervasive genome-wide negative correlation between promoter methylation and gene expression. If the negative correlation between DNA methylation level and expression level is directly involved in gene silencing in XCI, the female X chromosome would likely exhibit hypermethylation to silence the genes on the inactivated X chromosome. The result implies that the genome-wide pattern of negative correlation does not play a direct role in XCI in tree shrews. Future studies are needed to investigate other regulatory mechanisms that regulate XCI in this species.

While the global pattern of differential DNA methylation was not associated with differential gene expression of male and female X chromosome, we find evidence supporting the role of DNA methylation in the regulation of *Xist*, the key regulator of XCI in other mammals. We annotated *Xist* from the tree shrew genome, and show that the female and male X chromosomes generate distinctive transcripts. The transcripts from the male X chromosomes tended to be short and only a few copies were identified, which may potentially arise from unstable RNAs known to continue being expressed on the active X chromosome while stable RNAs are accumulating on the inactive X chromosome (Panning et al., 1997; Sheardown et al., 1997). The *Xist* was highly expressed in the female X chromosome and was strongly differentially methylated between the female and male X chromosomes. We further identified a regulatory region (CpG island) in the 5’ region of the *Xist* gene. The hypomethylation of this CpG island was tightly correlated with the expression of *Xist*. Moreover, hypomethylation of that specific CpG island was observed in other mammalian species. These observations indicate that DNA methylation is a key mechanism of regulation of *Xist*, which is the initiator of XCI.

We also explored DNA methylation of the Y-linked segments, utilizing the abundance of reads thanks to the next-generation sequencing approach. Epigenetics of the mammalian Y chromosome is currently little understood in large part due to the difficulties associated with the sequencing and assembly of the Y chromosome. Nevertheless, we show that contigs that could be best explained by the Y-linkage exhibit markedly reduced levels of DNA methylation compared to the X chromosome and autosomes (Fig. 5). Even though we have not and cannot attempt to identify specific genomic regions from these data, it is notable that DNA methylation levels of the same putative Y-contigs from the three males are highly correlated (Fig. 5), indicating that we are observing reproducible patterns of DNA methylation from the tree shrew Y-chromosome. The observation that the putative Y-linked segments exhibit reduced methylation supports the idea that DNA methylation tends to be reduced in less transcriptionally active regions (Makova et al., 2024) The comprehensive DNA methylome and gene expression data from tree shrews provide new insights into the evolution of genome, methylome and their interactions on the regulation of X chromosomes.

## Materials and Methods

### Tissue samples

The adult Chinese tree shrews used in this study were obtained from the Laboratory Animal Center of Kunming Institute of Zoology and handled in accordance with the guidelines approved by the Animal Care and Use Committee of the Kunming Institute of Zoology, Chinese Academy of Sciences. Following sacrifice of both male and female tree shrews, their brains were dissected and cortex tissues were rapidly frozen using liquid nitrogen for long-term storage at – 80 °C. All protocols of this study were approved by the internal review board of Kunming Institute of Zoology, Chinese Academy of Sciences (Sample information, Table S1).

### RNA sequencing

Total RNA was extracted from PFC tissues using TRIzol Reagent, and the integrity of RNA was assessed using Agilent 2100 bioanalyzer. The library preparation began with total RNA as the initial template. Specifically, mRNA with PolyA tails was enriched from total RNA using Oligo(dT) magnetic beads. The resulting mRNA was then randomly fragmented by divalent cations in fragmentation buffer. The first strand cDNA was synthesized in the M-MuLV reverse transcriptase system using the fragmented mRNA. The second strand cDNA was subsequently synthesized in the DNA polymerase I system using dNTPs. The obtained double-stranded cDNA was purified, end repaired, and then poly-A tails and sequencing adaptors were ligated. AMPure XP beads were used to screen for cDNA fragments within the size range of 370-420bp followed by PCR amplification and product purification. Library quality assessment was performed on the Agilent Bioanalyzer 2100 system, while cluster generation took place on the cBot Cluster Generation System. Finally, library preparations were sequenced on an Illumina Novaseq platform generating paired-end reads of 150 bp each.

### Whole-genome bisulfite sequencing

Genomic DNA was extracted from PFC tissues by phenol-chloroform extraction and ethanol precipitation. The extracted genomic DNA was quantified using Qubit fluorometer, and the 200ng of gDNA containing 1% unmethylated Lambda DNA was then randomly fragmented into 300bp small fragments using the Covaris LE220R ultrasonic fragmentation instrument. These small fragments were then subjected to terminal repair and adenylation before being fitted with methylated adapters. The bisulfite treatment step was performed using the EZ DNA Methylation-Gold kit (Zymo Research) following the manufacturer’s instructions. The resulting single-stranded DNA was PCR amplified and the PCR products were purified. Similarly, libraries obtained were quantified using Qubit fluorometer and their size distribution was analyzed by Agilent BioAnalyzer (Agilent). Paired-end sequencing was performed using an Illumina NovaSeq6000 according to Illumina-provided protocols. Finally, standardized WGBS data analysis pipeline was employed for analyzing the resulting data.

### Processing whole-genome bisulfite sequencing data

We first performed quality and adapter trimming using TrimGalore v0.6.7 (Babraham Institute) with paired-end mode and default parameters. Subsequently, reads were mapped to the tree shrew reference genome (TS_3.0) from the tree shrewDB (Fan et al., 2014; Ye et al., 2021), using Bismark v0.24.0 (Krueger & Andrews, 2011). Following deduplication using Bismark, we obtained coverage for over 96% CpG sites, with a read-depth between 12X-18X (Table. S1). The female4 and male4 samples exhibited relatively lower mapping efficiency. Additionally, we mapped the reads to the lambda phage genome (NC_001416.1) to estimate the bisulfite conversion rate in each sample, resulting in values 99.1-99.3% (Table S1).

The data, comprising the counts of methylated and unmethylated cytosines in each C-context at individual cytosines, were generated as cytosine report files using Bismark methylation extractor with –-bedGraph –-cytosine_report –-CX_context options. These output cytosine reports were used as input files for ViewBS v0.1.11 (Huang et al., 2018) to estimate global methylation levels and assess methylation patterns near gene and promoter regions. CpG sites that are detected as a positional difference of 1 and are located on different DNA strands are merged into a single CpG site, considering the symmetrical nature of CpG methylation. Subsequently, we calculated the fractional methylation level for each cytosine with at least five read counts by taking a ratio of methylated cytosine reads to the total read count. These outputs were employed in downstream analysis.

### Processing RNA-seq data

We mapped the RNA-seq reads to the tree shrew reference genome (TS_3.0) using HISAT2 v2.2.1 (Kim et al., 2019) with the –-dta option. Here the reference annotation information (Ye et al., 2021) were embedded in the genome index using HISAT2 –ss and –exon options. Transcripts were then assembled for each sample using StringTie v2.2.1 (Pertea et al., 2015), and we generated an updated GTF annotation included novel transcripts using the –merge flag. This process was guided by the reference GTF annotation using the StringTie –G flag. We obtained transcript and gene abundance information using the –eB and –A options. These output files were used to define gene regions and estimate expression levels.

Among the 224,473 transcripts identified by StringTie, 183,669 transcripts were guided by the reference genome, while the rest 40,804 transcripts were detected as novel transcripts, with 18,623 of them lacking strand information. In the downstream analysis, we excluded some transcripts generated from StringTie that lacked strand information for promoter methylation level calculation.

### Quantifying methylation levels in global and in gene bodies or promoter regions

We employed ViewBS v0.1.11 (Huang et al., 2018) to estimate the global weighted methylation levels and to analyze methylation landscape near gene and promoter regions. Cytosines in all C-context from all 30 chromosome and X chromosome were included in the analysis. We designated the longest transcript of each gene as a gene body in the process. For further analysis, we estimated the mean methylation level in the gene body region for each individual gene, including those with more than 3 CpG fractional methylation level values available. Promoter regions are defined as the 2kb upstream region of each gene body’s start site, and mean methylation levels are calculated in the same way.

### Identification of sex-specifically expressed gene

Following the processing of RNA-seq data, we obtained read count information for each sample. For the sex-specific gene analysis, we employed DESeq2 v1.38.3 (Love et al., 2014). We restricted our analysis to genes with at least 5 counts in 3 or more samples among total six samples (Fig. S9). We identified sex-specific genes with an adjusted p-value less than 0.1.

### Sex-specific read-depth on the X chromosome

Comparing the sex-specific read-depth on the X chromosome, the average read-depth in CpG sites were roughly twice in females compared to males for the X-linked sites, corresponding to the number of the X chromosome in females and males (Fig. S3, Fig. S7). We discovered that certain X-linked regions exhibited similar coverages between the female and male samples, which are potentially originated from the PAR (pseudoautosomal region) of the X chromosome. Intriguingly, we also observed one region on the X chromosome (Genomic coordinate 4542400-6144400) where female sample showed a complete lack of read counts (Fig. S3). We hypothesized that region may contain portion of the Y chromosome that was wrongly annotated. This region also harbored several potential Y chromosome genes that exhibited 0 expression in female samples but significant expression in male samples (Fig. S3). Based on these findings, we have concluded that this region is wrongly included in X chromosome reference from the Y chromosome. To address this issue, we have excluded both the CpG sites and genes located within this region. A total of 74 genes were identified within this region and subsequently removed in the X chromosome analysis. We identified an additional region with this characteristic on chromosome 26 (Genomic coordinate < 3132930) and excluded 85 genes in the region in sex-specific gene analysis.

### Identification of the *Xist* gene in the tree shrew genome

The *Xist* gene was not annotated in the reference annotation data. We initiated a process to locate the gene in the tree shrew X chromosome. Our approach involved performing BLASTN v2.13.0+ (Camacho et al., 2009) analysis using the human *Xist* gene (NR_001564.2) against the tree shrew reference genome. This allowed us to identify a potential range of the genomic coordinate of the *Xist* gene. Subsequently, we employed StringTie’s functionality for de novo transcript assembly (Pertea et al., 2015). We investigated the transcripts identified by StringTie and reference annotation in the potential range of the *Xist* gene. We found these transcripts exhibited female-specific expression patterns. Based on these findings, we defined these specific transcripts and their associated gene as the *Xist* gene located within the region of chrX 54,721,977-54,759,147 in the tree shrew genome. In examining differential DNA methylation between females and males within *Xist* region, we included all CpG sites detected at least twice in females or males and its fractional methylation levels were averaged in females and males each.

### Comparative studies among tree shrew, human, and mouse

To compare the sex-specific pattern near the *Xist* gene region within tree shrew, human, and mouse, we downloaded the public data for two male and two female samples of mouse fatal brains (Islam et al., 2022) from the Gene Expression Omnibus database with accession# GSE157553 and the data for one male and one female human sample of prefrontal cortex (Zeng et al., 2012) with accession# GSE37202. The processed files, along with the CpG coverage files for mice and the fractional methylation value files for humans, are formatted uniformly. CpG sites differing by a single position on different DNA strands are merged into one CpG site, aligning with the approach employed for tree shrews. Utilizing CpG sites with a read depth of 3 or greater, DNA methylation levels were calculated across 2kb-sized windows overlapping by 1kb around the *Xist* gene region for all species.

To compare the CpG density in the genomes of human, koala, and tree shrew, we calculated the CpG Observed/Expected ratio across 1000bp-sized windows (with 500bp overlaps) in the genome of each species. Each window is annotated as either a promoter (2000bp upstream of TSS) or gene body, depending on whether the window center falls within the promoter or gene body region. We utilized the human genome T2T-chm13v2.0 genome and gene annotation and the koala phaCin_unsw_v4.1 genome and RefSeq annotation for the analysis. To identify X-linked regions, we referenced the list of X-linked scaffolds provided in (Singh et al., 2021).

### Detection of Y-linked contigs and DNA methylation

We employed a bioinformatic method to identify putative Y-linked regions in our data. As the current tree shrew genome assembly does not provide a Y chromosome, we utilized the currently unassembled contigs in the tree shrew reference genome. We calculated the read-depth of cytosines in the CG context for each of these contigs and then averaged the values separately for male and female samples. Subsequently, for contigs containing more than 40 cytosines, we computed [1 – (mean read-depth in females)/(mean read-depth in males)]. Using this calculated metric distribution (Fig. 5), we generated a set of potentially 958 autosome-linked contigs (with values between –0.50 and 0.75), 563 X-linked contigs (with values smaller than – 0.5), and 93 Y-linked contigs (with values greater than 0.75). To validate our approach, we calculated the mean methylation levels for each set of contig and compared them. BLASTN v2.13.0+ (Camacho et al., 2009) is utilized to identify the sex-determining region Y (*Sry*)-related region in the tree shrew genome, employing the human *Sry* gene (NIH Gene ID 6736) and the rhesus monkey *Sry* gene (NIH Gene ID 574155).

## Author contributions

Lei Shi and Yifan Kong conceived the methylome and transcriptome sequencing. Yifan Kong, Yulian Tan and Ting Hu performed tree shrew brain dissection, RNA and DNA extraction, and generated the sequencing libraries. Dongmin Son performed the processing and computational analysis of genomic data, generated figures, and prepared the manuscript. Soojin Yi directed the analysis of the data, participated in developing analysis methods, interpreting the results, and preparing the manuscript. Lei Shi participated in preparing the manuscript and interpreting the results.

## Literature Cited

1. Al Adhami, H., Bardet, A. F., Dumas, M., Cleroux, E., Guibert, S., Fauque, P., Acloque, H., & Weber, M. (2022). A comparative methylome analysis reveals conservation and divergence of DNA methylation patterns and functions in vertebrates. BMC Biology, 20(1), 70. 10.1186/s12915-022-01270-x

2. Beard, C., Li, E., & Jaenisch, R. (1995). Loss of methylation activates Xist in somatic but not in embryonic cells. Genes & Development, 9(19), 2325–2334. 10.1101/gad.9.19.2325

3. Brown, C. J., Hendrich, B. D., Rupert, J. L., Lafrenière, R. G., Xing, Y., Lawrence, J., & Willard, H. F. (1992). The human XIST gene: Analysis of a 17 kb inactive X-specific RNA that contains conserved repeats and is highly localized within the nucleus. Cell, 71(3), 527–542. 10.1016/0092-8674(92)90520-m

4. Camacho, C., Coulouris, G., Avagyan, V., Ma, N., Papadopoulos, J., Bealer, K., & Madden, T. L. (2009). BLAST+: Architecture and applications. BMC Bioinformatics, 10(1), 421. 10.1186/1471-2105-10-421

5. Elango, N., & Yi, S. V. (2008). DNA Methylation and Structural and Functional Bimodality of Vertebrate Promoters. Molecular Biology and Evolution, 25(8), 1602–1608. 10.1093/molbev/msn110

6. Fan, Y., Huang, Z.-Y., Cao, C.-C., Chen, C.-S., Chen, Y.-X., Fan, D.-D., He, J., Hou, H.-L., Hu, L., Hu, X.-T., Jiang, X.-T., Lai, R., Lang, Y.-S., Liang, B., Liao, S.-G., Mu, D., Ma, Y.-Y., Niu, Y.-Y., Sun, X.-Q., … Yao, Y.-G. (2013). Genome of the Chinese tree shrew. Nature Communications, 4(1), 1426. 10.1038/ncomms2416

7. Fan, Y., Yu, D., & Yao, Y.-G. (2014). Tree shrew database (TreeshrewDB): A genomic knowledge base for the Chinese tree shrew. Scientific Reports, 4(1), Article 1. 10.1038/srep07145

8. Gimelli, G., Giorda, R., Beri, S., Gimelli, S., & Zuffardi, O. (2006). A 46,X,inv(Y) young woman with gonadal dysgenesis and gonadoblastoma: Cytogenetics, molecular, and methylation studies. American Journal of Medical Genetics. Part A, 140(1), 40–45. 10.1002/ajmg.a.31044

9. Hellman, A., & Chess, A. (2007). Gene body-specific methylation on the active X chromosome. Science (New York, N.Y.), 315(5815), 1141–1143. 10.1126/science.1136352

10. Hendrich, B. D., Brown, C. J., & Willard, H. F. (1993). Evolutionary conservation of possible functional domains of the human and murine XIST genes. Human Molecular Genetics, 2(6), 663–672. 10.1093/hmg/2.6.663

11. Huang, X., Zhang, S., Li, K., Thimmapuram, J., Xie, S., & Wren, J. (2018). ViewBS: A powerful toolkit for visualization of high-throughput bisulfite sequencing data. Bioinformatics (Oxford, England), 34(4), 708–709. 10.1093/bioinformatics/btx633

12. Islam, M., Strawn, M., & Behura, S. K. (2022). Fetal origin of sex-bias brain aging. FASEB Journal: Official Publication of the Federation of American Societies for Experimental Biology, 36(8), e22463. 10.1096/fj.202200255RR

13. Jeong, H., Mendizabal, I., Berto, S., Chatterjee, P., Layman, T., Usui, N., Toriumi, K., Douglas, C., Singh, D., Huh, I., Preuss, T. M., Konopka, G., & Yi, S. V. (2021). Evolution of DNA methylation in the human brain. Nature Communications, 12(1), 2021. 10.1038/s41467-021-21917-7

14. Jjingo, D., Conley, A. B., Yi, S. V., Lunyak, V. V., & Jordan, I. K. (2012). On the presence and role of human gene-body DNA methylation. Oncotarget, 3(4), 462–474. 10.18632/oncotarget.497

15. Keown, C. L., Berletch, J. B., Castanon, R., Nery, J. R., Disteche, C. M., Ecker, J. R., & Mukamel, E. A. (2017). Allele-specific non-CG DNA methylation marks domains of active chromatin in female mouse brain. Proceedings of the National Academy of Sciences of the United States of America, 114(14), E2882–E2890. 10.1073/pnas.1611905114

16. Kim, D., Paggi, J. M., Park, C., Bennett, C., & Salzberg, S. L. (2019). Graph-based genome alignment and genotyping with HISAT2 and HISAT-genotype. Nature Biotechnology, 37(8), Article 8. 10.1038/s41587-019-0201-4

17. Krueger, F., & Andrews, S. R. (2011). Bismark: A flexible aligner and methylation caller for Bisulfite-Seq applications. Bioinformatics, 27(11), 1571–1572. 10.1093/bioinformatics/btr167

18. Larney, C., Bailey, T. L., & Koopman, P. (2014). Switching on sex: Transcriptional regulation of the testis-determining gene Sry. Development (Cambridge, England), 141(11), 2195–2205. 10.1242/dev.107052

19. Li, H., Xiang, B.-L., Li, X., Li, C., Li, Y., Miao, Y., Ma, G.-L., Ma, Y.-H., Chen, J.-Q., Zhang, Q.-Y., Lv, L.-B., Zheng, P., Bi, R., & Yao, Y.-G. (2024). Cognitive Deficits and Alzheimer’s Disease-Like Pathologies in the Aged Chinese Tree Shrew. Molecular Neurobiology, 61(4), 1892–1906. 10.1007/s12035-023-03663-7

20. Lister, R., Pelizzola, M., Dowen, R. H., Hawkins, R. D., Hon, G., Tonti-Filippini, J., Nery, J. R., Lee, L., Ye, Z., Ngo, Q.-M., Edsall, L., Antosiewicz-Bourget, J., Stewart, R., Ruotti, V., Millar, A. H., Thomson, J. A., Ren, B., & Ecker, J. R. (2009). Human DNA methylomes at base resolution show widespread epigenomic differences. Nature, 462(7271), 315–322. 10.1038/nature08514

21. Love, M. I., Huber, W., & Anders, S. (2014). Moderated estimation of fold change and dispersion for RNA-seq data with DESeq2. Genome Biology, 15(12), 550. 10.1186/s13059-014-0550-8

22. Makova, K. D., Pickett, B. D., Harris, R. S., Hartley, G. A., Cechova, M., Pal, K., Nurk, S., Yoo, D., Li, Q., Hebbar, P., McGrath, B. C., Antonacci, F., Aubel, M., Biddanda, A., Borchers, M., Bornberg-Bauer, E., Bouffard, G. G., Brooks, S. Y., Carbone, L., … Phillippy, A. M. (2024). The complete sequence and comparative analysis of ape sex chromosomes. Nature, 1–11. 10.1038/s41586-024-07473-2

23. Mendizabal, I., Berto, S., Usui, N., Toriumi, K., Chatterjee, P., Douglas, C., Huh, I., Jeong, H., Layman, T., Tamminga, C. A., Preuss, T. M., Konopka, G., & Yi, S. V. (2019). Cell type-specific epigenetic links to schizophrenia risk in the brain. Genome Biology, 20(1), 135. 10.1186/s13059-019-1747-7

24. Morgan, R., Loh, E., Singh, D., Mendizabal, I., & Yi, S. V. (2024). DNA methylation differences between the female and male X chromosomes in human brain (p. 2024.04.16.589778). bioRxiv. 10.1101/2024.04.16.589778

25. Nishino, K., Hattori, N., Tanaka, S., & Shiota, K. (2004). DNA methylation-mediated control of Sry gene expression in mouse gonadal development. The Journal of Biological Chemistry, 279(21), 22306– 22313. 10.1074/jbc.M309513200

26. Norris, D. P., Patel, D., Kay, G. F., Penny, G. D., Brockdorff, N., Sheardown, S. A., & Rastan, S. (1994). Evidence that random and imprinted Xist expression is controlled by preemptive methylation. Cell, 77(1), 41–51. 10.1016/0092-8674(94)90233-x

27. Panning, B., Dausman, J., & Jaenisch, R. (1997). X chromosome inactivation is mediated by Xist RNA stabilization. Cell, 90(5), 907–916. 10.1016/s0092-8674(00)80355-4

28. Panning, B., & Jaenisch, R. (1996). DNA hypomethylation can activate Xist expression and silence X-linked genes. Genes & Development, 10(16), 1991–2002. 10.1101/gad.10.16.1991

29. Penny, G. D., Kay, G. F., Sheardown, S. A., Rastan, S., & Brockdorff, N. (1996). Requirement for Xist in X chromosome inactivation. Nature, 379(6561), 131–137. 10.1038/379131a0

30. Pertea, M., Pertea, G. M., Antonescu, C. M., Chang, T.-C., Mendell, J. T., & Salzberg, S. L. (2015). StringTie enables improved reconstruction of a transcriptome from RNA-seq reads. Nature Biotechnology, 33(3), Article 3. 10.1038/nbt.3122

31. Pontier, D. B., & Gribnau, J. (2011). Xist regulation and function eXplored. Human Genetics, 130(2), 223– 236. 10.1007/s00439-011-1008-7

32. Rauch, T. A., Wu, X., Zhong, X., Riggs, A. D., & Pfeifer, G. P. (2009). A human B cell methylome at 100−base pair resolution. Proceedings of the National Academy of Sciences, 106(3), 671–678. 10.1073/pnas.0812399106

33. Schübeler, D. (2015). Function and information content of DNA methylation. Nature, 517(7534), Article 7534. 10.1038/nature14192

34. Sheardown, S. A., Duthie, S. M., Johnston, C. M., Newall, A. E., Formstone, E. J., Arkell, R. M., Nesterova, T. B., Alghisi, G. C., Rastan, S., & Brockdorff, N. (1997). Stabilization of Xist RNA mediates initiation of X chromosome inactivation. Cell, 91(1), 99–107. 10.1016/s0092-8674(01)80012-x

35. Singh, D., Sun, D., King, A. G., Alquezar-Planas, D. E., Johnson, R. N., Alvarez-Ponce, D., & Yi, S. V. (2021). Koala methylomes reveal divergent and conserved DNA methylation signatures of X chromosome regulation. Proceedings of the Royal Society B: Biological Sciences, 288(1945), 20202244. 10.1098/rspb.2020.2244

36. Sun, D., Maney, D. L., Layman, T. S., Chatterjee, P., & Yi, S. V. (2019). Regional epigenetic differentiation of the Z Chromosome between sexes in a female heterogametic system. Genome Research, 29(10), 1673–1684. 10.1101/gr.248641.119

37. Tinker, A. V., & Brown, C. J. (1998). Induction of XIST expression from the human active X chromosome in mouse/human somatic cell hybrids by DNA demethylation. Nucleic Acids Research, 26(12), 2935–2940. 10.1093/nar/26.12.2935

38. Yamashita, A., Fuchs, E., Taira, M., Yamamoto, T., & Hayashi, M. (2012). Somatostatin-immunoreactive senile plaque-like structures in the frontal cortex and nucleus accumbens of aged tree shrews and Japanese macaques. Journal of Medical Primatology, 41(3), 147–157. 10.1111/j.1600-0684.2012.00540.x

39. Yao, Y.-G. (2017). Creating animal models, why not use the Chinese tree shrew (Tupaia belangeri chinensis)? Zoological Research, 38(3), 118–126. 10.24272/j.issn.2095-8137.2017.032

40. Ye, M.-S., Zhang, J.-Y., Yu, D.-D., Xu, M., Xu, L., Lv, L.-B., Zhu, Q.-Y., Fan, Y., & Yao, Y.-G. (2021). Comprehensive annotation of the Chinese tree shrew genome by large-scale RNA sequencing and long-read isoform sequencing. Zoological Research, 42(6), 692–709. 10.24272/j.issn.2095-8137.2021.272

41. Zeng, J., Konopka, G., Hunt, B. G., Preuss, T. M., Geschwind, D., & Yi, S. V. (2012). Divergent whole-genome methylation maps of human and chimpanzee brains reveal epigenetic basis of human regulatory evolution. American Journal of Human Genetics, 91(3), 455–465. 10.1016/j.ajhg.2012.07.024

